# Constructing founder sets under allelic and non-allelic homologous recombination

**DOI:** 10.1101/2022.05.27.493721

**Authors:** Konstantinn Bonnet, Tobias Marschall, Daniel Doerr

## Abstract

Homologous recombination between the maternal and paternal copies of a chromosome is a key mechanism for human inheritance and shapes population genetic properties of our species. However, a similar mechanism can also act between different copies of the same sequence, then called *non-allelic homologous recombination* (NAHR). This process can result in genomic rearrangements—including deletion, duplication, and inversion—and is underlying many genomic disorders. Despite its importance for genome evolution and disease, there is a lack of computational models to study genomic loci prone to NAHR.

In this work, we propose such a computational model, providing a unified framework for both (allelic) homologous recombination and NAHR. Our model represents a set of genomes as a graph, where human haplotypes correspond to walks through this graph. We formulate two founder set problems under our recombination model, provide flow-based algorithms for their solution, and demonstrate scalability to problem instances arising in practice.

## 1 Introduction

Twenty years ago, at this conference^1^, Esko Ukkonen introduced the problem of inferring founder sets from haplotyped SNP sequences under allelic recombination [30]. Ukkonen’s work has since inspired a wealth of research addressing various aspects and applications of founder set reconstruction ranging from the reconstruction of ancestral recombinations and pangenomics to applications in phage evolution [16, 19, 29]. In its original setting, the problem sets out from a given set of *m* sequences of equal length *n*, where characters across sequences residing at the same index position correspond to a SNP. It then asks for a smallest set of sequences, called *founder set*, such that each given sequence can be constructed through a series of crossovers between sequences of the founder set, where each segment between two successive recombinations must meet a minimum length threshold. The *Founder Set Reconstruction* problem is NP-hard in general [22], but is solvable in linear time for the special case of founder sets of size two [30, 35]. Since its introduction, various heuristics and approximations have been proposed [35, 24, 25]. A variant of this problem restricts crossovers to coincide at certain positions, thereby decomposing the input sequences into a universally shared succession of blocks. The resulting problem, known as *Minimum Segmentation Problem* is polynomial [26]. In his seminal paper, Ukkonen devised a *O*(*n*^2^*m*) algorithm for its solution which has been improved by Norri *et al*. [17] to linear time, i.e. *O*(*nm*), capitalizing on recent breakthroughs in data structures [9].

Just like the Founder Set Problem, the vast majority of population genetic analyses and genomewide association studies have been focused on SNPs in the past decades, neglecting the more complex forms of variation—mostly for technical difficulties in detecting them. In particular, structural variants (SVs), commonly defined as variants of at least 50bp, have posed substantial challenges and studies based on short sequencing reads typically detect less than half of all SVs present in a genome [37]. Recent technological and algorithmic advances help to overcome these limitations [27]. Long read technologies now enable haplotype-resolved *de novo* assembly of human genomes [20], which in turn enables a much more complete ascertainment of SVs [10]. Earlier this year, the first complete telomere-to-telomere assembly of a human genome was announced [18], heralding a new era of genomics where high-quality, haplotype-resolved assemblies of complex repetitive genomic structures become broadly available. Presently, the Human Pangenome Reference Consortium (HPRC), is applying these techniques to generate a large panel of haplotype-resolved genome assemblies from samples of diverse ancestries [33]. These emerging data sets enable studying genetic loci involving duplicated sequence, called *segmental duplications* (SDs), which are amenable to NAHR and are therefore highly mutable and show complicated evolutionary trajectories [13, 31]. The T2T-CHM13 study alone reports over 40 thousand segmental duplications that amount to 202 Mb (6.6% of the human genome) [18].

Interestingly, at loci with highly similar segments arranged in opposite orientations, such as Segment 3 in Figure 1, NAHR can lead to *inversion*, i.e. the reversal of the interior sequence (Segment 4 in Figure 1). Because of being flanked by a pair of copies of the same sequence (cf. Segment 3) that often comprises tens of thousands of bases, such events have been largely undetectable by sequencing technologies with read lengths below the length of the duplicated sequence; in particular by conventional short read sequencing. Recent studies applying multiple technologies reveal that inversions affect tens of megabases of sequence in a typical human genome [7]. Unlike most other classes of genetic variation, inversions are often *recurrent* with high mutation rates, that is, the same events have happened multiple times in human history [21]. Depending on the structures of duplicated sequence at a particular locus, individual human haplotypes can differ in their potential for NAHR. This can have important implications for the risk for a range of genetic disorders caused by NAHR-mediated mutations [21].

**Figure 1:**
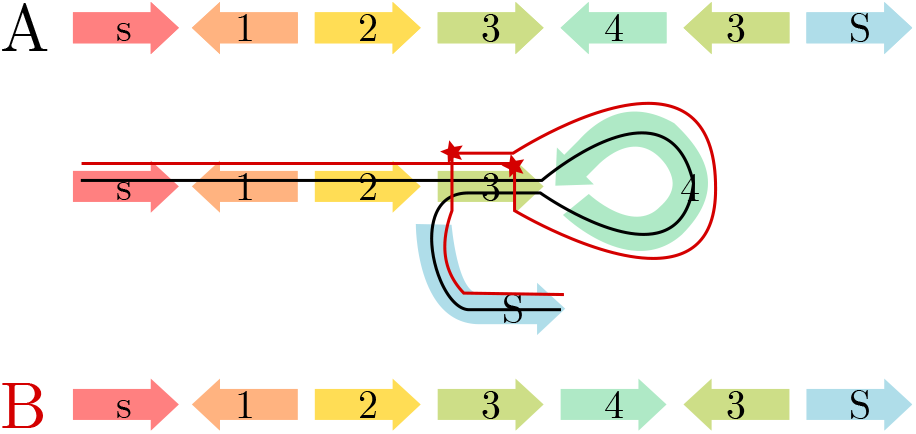
Illustration of an NAHR-mediated inversion. Haplotype *A* (black line) represents the original configuration, while haplotype *B* (red line) can be derived from *A* by two recombination events between inverted repeats of genomic marker 3 as indicated by the red stars.

In the past two decades, various mathematical models and algorithms to study genome rearrangements have been proposed. These range from the classic reversal [3, 2] and transposition [4] model to composed models for two or more balanced rearrangements [32, 8], to generalized models such as the popular *double cut and join* (DCJ) model [36, 5]. As the research in this field continues, advanced models can additionally accommodate one or more types of unbalanced rearrangements, i.e., deletion, insertion, and duplication [28, 6]. Yet, none of these models adequately considers sequence similarity as a prerequisite for NAHR, which is a key molecular mechanism shaping complex loci in the human genome. In summary, there are now technological opportunities to study the population history of recalcitrant SD loci that are prone to genome rearrangements and relevant to disease, but computational models to facilitate this have so far been lacking.

In this work, we study homologous recombination in a genome model that represents DNA sequences at a level of abstraction where they are already decomposed into genomic markers with assigned homologies. Here, our notion of homology is a synonym for *high DNA sequence similarity*, as we adopt the terminology underlying the concept of homologous recombination. Our model permits recombination events to occur between homologous markers independent of their position within or between haplotypes, as long as the markers’ orientations are respected. In other words, a marker can only recombine with a homologous marker alongside the same direction, as illustrated by Figure 1, because a recombination event can only occur between homologous markers if they are aligned to each other. By virtue of recapitulating the underlying molecular mechanism (NAHR), it implicitly allows for all the rearrangements it can give rise to, including deletion, duplication, and inversion.

Marker decomposition and homology assignment can be done in practice with genome graph builders such as MBG [23], minigraph [12], or pggb^2^. In fact, our algorithms are based on *variation graph* or *pangenome graph*, where nodes correspond to homologous DNA segments and edges between segments correspond to observed adjacencies in a given set of haplotypes.

## 2 Methods

### 2.1 Preliminaries

A *(genomic) marker m* is an element of the finite universe of markers denoted by ℳ, and is associated with a fragment of a double-stranded DNA molecule. Each marker can be traversed in *forward* and *reverse* direction. A marker in forward orientation (which is the default orientation) is traversed from left to right. Overline notation 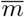 indicates the reversal of a marker *m*, which is carried out relative to its orientation, i.e., 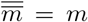. Similarly, 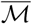 represents the set of all reverse oriented markers. We designate two forward oriented markers {*s, S*} ⊆ ℳ as *terminal markers*. In what follows, we study *terminal sequences*, that is, sequences drawn from the alphabet of oriented markers 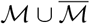 that start with *s* or 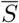, end in *S* or 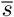 and do not contain any further terminal markers in between. A terminal sequence can be traversed in forward and reverse direction. A *haplotype* is a terminal sequence that starts with *s* (*source*) and ends with *S* (*sink*).

**Example 1**. *Consider in the following two sequences of genomic markers A and X drawn from the universe of markers* `ℳ = {s, 1, 2, 3, 4, S}`, *where* 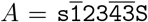 *and* 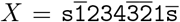. *Sequence A starts and ends with terminal markers* s *and* S, *respectively, thus constituting a* haplotype *drawn from* ℳ. *Conversely, X starts with* `s` *and ends in* 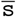 *and therefore is a terminal sequence, but not a* haplotype.

Given a sequence *A*, |*A*| indicates the length of *A* which corresponds to the number of *A*’s constituting elements. 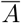 defines the *reverse complementation* of sequence *A*, i.e., the simultaneous reversal of the sequence and its constituting elements. The element at the *i*th position in sequence *A* is denoted by *A*[*i*]. A *segment* of sequence *A* starting at position *i* and ending at and including position *j*, is denoted by *A*[*i*..*j*]. In particular, *A*[..*i*] := *A*[1..*i*] and *A*[*i*..] := *A*[*i*..|*A*|] denote the *prefix* and *suffix* of *A*, respectively. The operator “+” indicates the concatenation of two sequences.

**Example 1** (cont’d). *The length of A is* |*A*| = 7; *its reverse complement is* 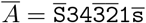; *A*[4..6] *is a segment of A and corresponds to sequence* 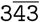; *The segments* 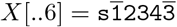 *and A*[7..] = S *are a prefix and a suffix of X and A, respectively; The concatenation of prefix X*[..6] *and suffix A*[7..] *results in haplotype* 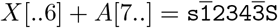.

A *recombination* is an operation that acts on a shared oriented marker *m* of any two terminal sequences *A* and *B*: let *A*[*i*] = *B*[*j*] = *m*; recombination *χ*(*A, B, i, j*) produces terminal sequence *C* = *A*[..*i*] + *B*[*j* + 1..]. For a given set of haplotypes ℋ, span(ℋ) denotes the *span*, i.e., the set of all *haplotypes* generated by applying *χ* on haplotypes ℋ and the resulting terminal sequences. More precisely, let 𝒯 be the universe of terminal sequences, defined recursively by 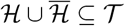 such that for any *A, B* ∈ 𝒯 with some *A*[*i*] = *B*[*j*] the recombinant *C* = *A*[*i*] + *B*[*j* + 1] and its reverse complement 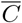 is also in 𝒯. Then *span*(ℋ) := {*A* ∈ 𝒯 | *A* is a haplotype}. Accordingly, we also say that “ℋ is a *generating set* of span(ℋ)”. Conversely, given any (possibly infinite) set of haplotypes 𝒮 and some ℋ ⊆ 𝒮, ℋ is a generating set of 𝒮 iff span(ℋ) = 𝒮.

**Example 1** (cont’d). *Recombination* 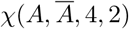 *produces terminal sequence* 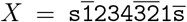. *Subsequent recombination χ*(*X, A*, 6, 6), *produces haplotype* 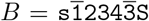. *If* {*A*} *is a given set of haplotypes, then* span({*A*}) = {*A, B*}.

In this paper, we study the following two problems:

**Problem 1** (Founder Set). *Given a set of haplotypes* ℋ, *find a generating set* ℱ ⊆ span(ℋ) *such that* Σ_*A*∈ℱ_ |*A*| *is minimized*.

We call a solution to Problem 1 a *founder set* and its members *founder sequences*.

**Problem 2**. *Given a set of haplotypes* ℋ, *find a founder set* ℱ *that minimizes the number of recombinations applied to haplotypes* ℋ *and their intermediate terminal sequences in constructing* ℱ.

### 2.2 Constructing Founder Sets

#### Variation graph construction

We solve Problem 1 by studying the *variation graph* 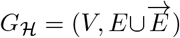 of the given set of haplotypes ℋ. Graph *G*_ℋ_ is an undirected edge-colored multigraph where each edge can have one of two colors corresponding to their membership in edge sets *E* and 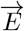. In constructing *G*_ℋ_, each marker *m* of the universe of forward-oriented markers ℳ is represented by a tuple of its *extremities* (*m*^t^, *m*^h^)^3^ also called “*tail*” and “*head*” of *m*, respectively, and its reverse-oriented counterpart 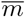 is represented as (*m*^h^, *m*^t^)^3^. Node set *V* of graph *G*_ℋ_ corresponds to the set of all marker extremities, and each marker *m* ∈ ℳ gives rise to one *marker edge* 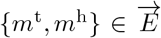. Further, any two (not necessarily distinct) nodes 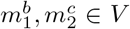 are connected by one *adjacency edge* 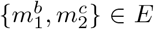 iff there exists a sequence 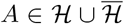 with *A* = ..*m*_1_*m*_2_.. such that 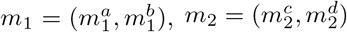,and {*a, b*} = {*c, d*} = {t, h}.

**Example 2**. *Let* 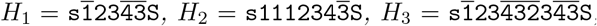, *and* 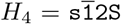, *then the variation graph G*_ℋ_ *of* ℋ= {*H*_1_, *H*_2_, *H*_3_, *H*_4_} *is as follows, with marker edges drawn in gray and adjacency edges in black:*

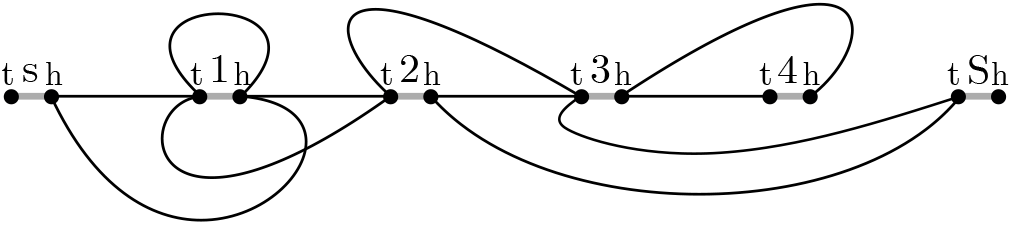

##### Proposition 1.

*Let G*_ℋ_ *be the variation graph of haplotypes* ℋ, *and χ the set of all walks between terminal markers s*^t^ *and S*^h^ *in G*_ℋ_ *with edges alternating between E and* 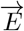, *then* span(ℋ) = *χ*.

*Proof*.

⇒ Observe that no recombination can create a new pair of consecutive markers *m*_1_*m*_2_ that is not contained in any sequence 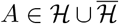. Therefore, each haplotype *B* ∈ span(ℋ) is a succession of consecutive markers drawn from sequences in 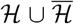, i.e., *B* can be delineated in *G*_ℋ_ by following adjacency edges corresponding to its succession of consecutive markers.
⇐ If each alternating walk *X* = (*s*^t^, *s*^h^, …, *S*^t^, *S*^h^) ∈ *χ* in variation graph *G*_ℋ_ corresponds to a haplotype *B* ∈ span(ℋ), then *X* must be producible through a series of recombinations of haplotypes H and their recombinants. We show this by construction:
  a. Pick some haplotype *A* ∈ ℋ and initialize *i* ← 1;
  b. Let 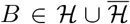 be a sequence such that for some position *j, B*[*j*..*j* + 1] = *m*_1_*m*_2_ with *m*_1_ = *X*[*i*..*i* + 1] and *m*_2_ = *X*[*i* + 2..*i* + 3]. Then *A* ← *χ*(*A, B, i/*2, *j*).
  c. Increase *i* by 2 and repeat step **b** unless *i* = |*X*| − 3.

Observe that by construction of the variation graph *G*_ℋ_, a suitable sequence 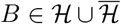 must exist in each iteration of step **b**.

#### Defining flows on variation graphs

We determine a minimum set of founder sequences by solving a network flow problem in variation graph *G*_ℋ_ where flow is allowed to travel along adjacency edges in either direction. In doing so, we find a non-negative flow *ϕ* : *V* × *V* → ℕ such that the total flow Σ_*u,v*∈*V*_*ϕ*(*u, v*) of graph *G*_ℋ_ is minimized and satisfies the following constraints:

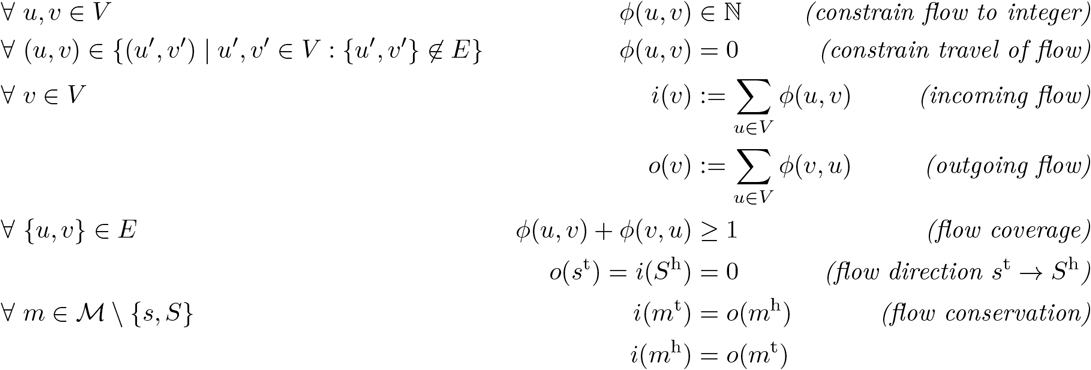

Note that the flow can travel in both directions of an edge {*u, v*} ∈ *E* and that *ϕ*(*u, v*) = *ϕ*(*v, u*) does not hold true in general. The only node pairs of the graph that are *unbalanced*, i.e., do not satisfy flow conservation, are (*s*^t^, *s*^h^) and (*S*^t^, *S*^h^).

**Example 2** (cont’d). *The drawing below illustrates a flow solution on variation graph G*_ℋ_, *with the direction and amount of flow along adjacency edges indicated by labeled arrowed arcs*.

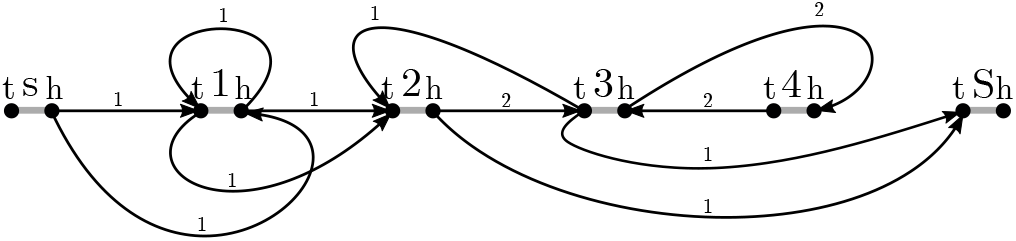

#### Deriving haplotypes from flows

By applying the Flow Decomposition Theorem [1, p. 80f], any *flow*, i.e., solution to the above-specified constraints, is decomposable into a set of alternating paths going from source *s*^t^ to sink *S*^h^ and a set of alternating cycles. Ahuja *et al*. [1] give a simple and efficient algorithm that does so in polynomial time and which we describe below, adapted to our circumstances. The idea is to perform a random walk in the graph from source to sink or within a cycle, thereby consuming flow along adjacency edges until all flow is depleted. The proof of the algorithm remains unchanged to that given by Ahuja *et al*., thus is not repeated here.

1. Set *u* ← *s*^t^.
2. Setting out from current node *u*, traverse the incident marker edge to some node *v*, choose any neighbor *w* of *v* for which *ϕ*(*v, w*) *>* 1. Follow the adjacency edge to *v* and decrease the flow *ϕ*(*v, w*) by 1. Set *u* ← *w*.
3. As long as *u* ≠ *S*^t^ do as follows: if *u* has been visited in the traversal before, then extract the corresponding alternating cycle from the recorded sequence and report it. Proceed with the traversal by repeating step 2.
4. However, if *u* = *S*^t^, follow the marker edge to *S*^h^ and report the recorded sequence as a path.
5. If *s*^h^ is incident to edges with positive flow, proceed with step 1. Otherwise, there still might be strictly positive flow remaining in the graph corresponding to unreported cycles. In that case, pick any node *u* ← *m*^*a*^ such that for some node *w, ϕ*(*m*^*b*^, *w*) *>* 0, {*a, b*} = {t, h} and *m* ∈ ℳ, and proceed with step 2.

**Example 2** (cont’d). *The components of the flow solution on variation graph G*_ℋ_ *comprise two cycles C1 and C2, and two* (*s*^*t*^, *S*^*h*^)*-paths P1 and P2, as illustrated below*.

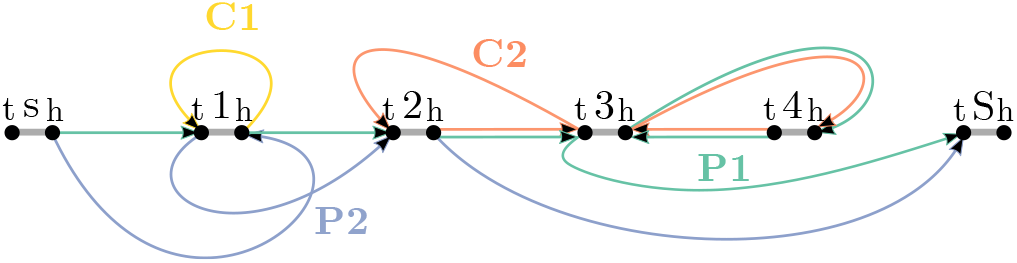

What remains is the integration of cycles into walks that then correspond to the haplotypes of the founder set. The integration is facilitated by a graph structure, the *component graph*. The component graph *G′* = (*V′, E′, l*) is an edge-labeled, directed multigraph, where, in its initial construction, each alternating (*s*^t^, *S*^h^)-path and each cycle reported during flow decomposition is represented by a distinct node of *V′*. In the component graph *G′*, each cycle *c* of the flow decomposition sharing one or more markers with another component *c′* is connected by one or more directed edges (*c, c′*) to that component, with each edge’s label *l*(*c, c′*) corresponding to one distinct shared marker, oriented according to the their succession in *c* (which may not be the same as in *c′*). The component graph is then successively deconstructed until empty as follows:

1. Remove and report all (*s*^t^, *S*^h^)-walks with in-degree 0 from node set *V′*^4^.
2. Pick a cycle *c* ∈ *V′* with in-degree 0, or, if none such exists, any arbitrary cycle *c* ∈ *V′*.
3. Pick an outgoing edge (*c, c′*) ∈ *E′* such that *c′* is a (*s*^t^, *S*^h^)-walk. If no such *c′* exists, *c* is only adjacent to cycles, out of which one *c′* is picked at will. Let (*m*^*a*^, *m*^*b*^) ← *l*(*c, c′*), {*a, b*} = {t, h}. If marker *m* is embedded in *c′* in same orientation, i.e. *c′* = ..*m*^*a*^*m*^*b*^.., then linearize *c* in *m*, i.e., *c* = *m*^*b*^*c*_1_..*c*_*k*−1_*m*^*a*^, and integrate it into *c′* such that *c′* ←..*m*^*a*^*m*^*b*^*c*_1_..*c*_*k*−1_*m*^*a*^*m*^*b*^... Otherwise, integrate the reversed linearization of *c*, i.e, *c′* ←..*m*^*b*^*m*^*a*^*c*_*k-*1_..*c*_1_*m*^*b*^*m*^*a*^... Remove cycle *c* and its outgoing edges from component graph *G*^*′*^.
4. Proceed with step 1 until no more components remain and all (*s*^t^, *S*^h^)-walks are reported.

The search for components with in-degree 0 can be efficiently implemented through preorder traversal of *G*^*′*^. Note that each cycle must have at least one outgoing edge and that ultimately all cycles *must be* integrable into a (*s*^t^, *S*^h^)-walk, otherwise this would imply that *G*_ℋ_ contains a disconnected, circular component that is not reachable by an alternating path from source *s*^t^ to sink *S*^h^, thus contradicting the correctness of *G*_ℋ_’s construction. The reported (*s*^t^, *S*^h^)-walks represent the wanted haplotypes of the founder set.

**Example 2** (cont’d). *The plot below depicts the component graph of components C1, C2, P1, and P2 (left) and the final two* (*s*^*t*^, *S*^*h*^)*-walks that collectively represent a founder set of* ℋ *(right)*.

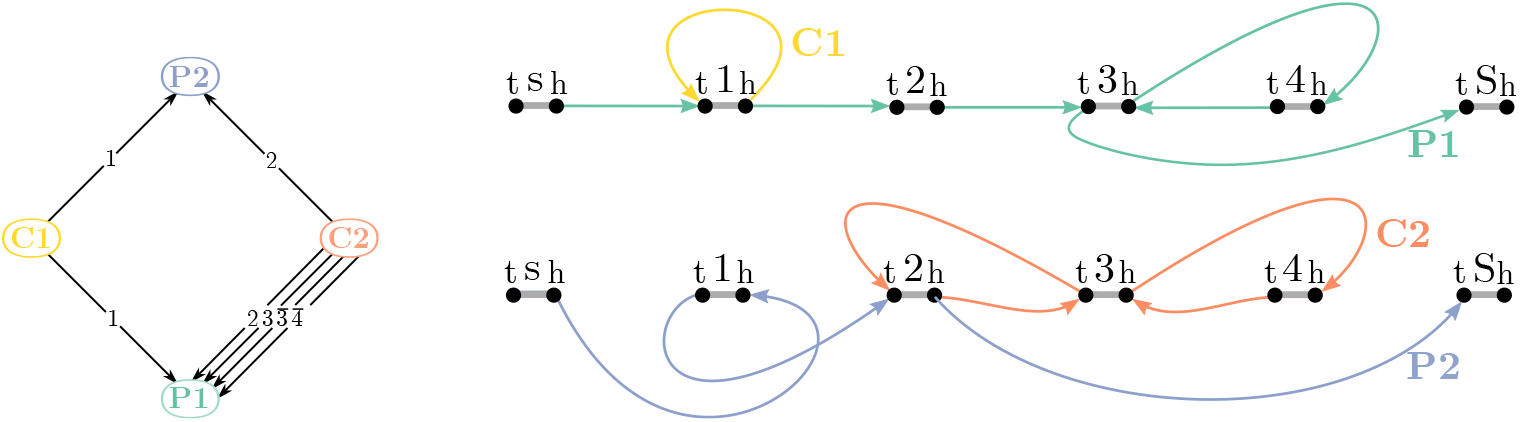

##### Theorem 1.

*Any flow that minimizes the total flow* Σ_*u,v*∈*V*_ *ϕ*(*u, v*) *of variation graph* 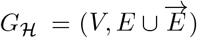 *of a given set of haplotypes* ℋ *is equivalent to a solution to Problem 1*.

*Proof*. It is sufficient to show that every flow is decomposable into a set of haplotypes (⇒) and every founder set represents a valid flow (⇐).

⇒ Any flow of variation graph *G*_H_ is decomposable into a set of haplotypes ℋ*′*, as demonstrated above. Observe that the above-listed flow constraints enforce the derived haplotypes ℋ*′* to cover the entire graph *G*_ℋ_ and consequently *G*_ℋ′_ = *G*_ℋ_. This implies that span(ℋ′) = span(ℋ), i.e., ℋ′ is a generating set of span(ℋ). Therefore, the sum of lengths of haplotypes derived from a flow solution is an upper bound of Problem 1.
⇐ Any set of haplotypes H*′* ⊆ span(ℋ) that covers each consecutive pair of markers *m*_1_*m*_2_ in haplotypes ℋ at least once (either in forward orientation *m*_1_*m*_2_ or in reverse orientation 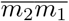) represents a valid flow of *G*_ℋ_. To construct a flow from ℋ′, set 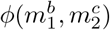 to the number of occurrences of consecutive markers *m*_1_*m*_2_ in haplotypes of ℋ′ with 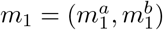 and 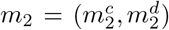, {*a, b*} = {*c, d*} = {t, h}. Observe that by construction, flow is integer, travels from source *s*^t^ to sink *S*^h^ and satisfies coverage and conservation constraints.

### 2.3 Minimizing Recombinations in Founder Sequences

We now present an algorithm towards solving Problem 2, i.e., the problem of finding a founder set that minimizes the number of recombinations needed for its construction from a given set of haplotypes ℋ. Our approach is exact under the assumption that the overall multiplicity of each pair of consecutive markers in the founder set of a solution to Problem 2 is known, yet the pair’s particular orientation in a founder sequence may be unresolved. To this end, we presume a given function 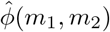 acting as oracle for the overall multiplicity of any given pair of consecutive oriented markers 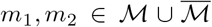^5^. More specifically, 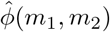 reports the total number of occurrences of *m*_1_*m*_2_ and 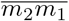 in a solution to Problem 2. In addition, we make use of function 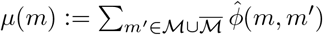 to retrieve the multiplicity of any marker 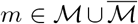^6^. Our solution makes use of the *flow graph* that is defined in the subsequent paragraph. We calculate a matching in the flow graph that describes a set of founder sequences, each corresponding to a succession of segments of haplotypes ℋ. The objective of the matching is to minimize the total number of these segments across all founder sequences which is equivalent to minimizing the number of recombinations for their construction from haplotype set ℋ.

#### Flow graph construction

The flow graph 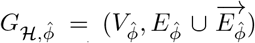 is a directed edge-colored multigraph with adjacency edges 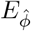 and marker edges 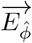, where each marker extremity *m*^*a*^, *m* ∈ ℳ and *a* ∈ {t, h}, gives rise to 2 · *μ*(*m*) elements in node set 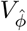, representing *μ*(*m*) many “*in*” (*i*) and *μ*(*m*) many “*out*” (*o*) nodes. That is, 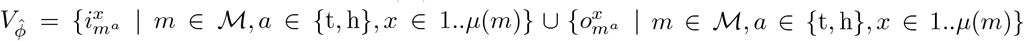. Each out node 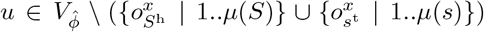 is incident to *one and only one* directed adjacency edge (*u, v*) connecting *u* to some in node *v* thereby realizing one occurrence of its representing pair of consecutive oriented markers in a founder sequence. Conversely, each forward-oriented marker *m* ∈ ℳ contributes *μ*(*m*)^2^ many directed marker edges that connect in/tail nodes with out/head nodes, i.e., 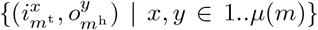. Analogously, each reverse-oriented marker 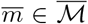 contributes *μ*(*m*)^2^ many in/head-to-out/tail-directed marker edges 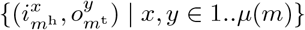.

**Example 2** (cont’d). *The flow graph* 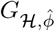 *for the given set of haplotypes* 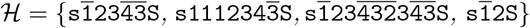 *and a given* 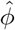 *is as follows:*

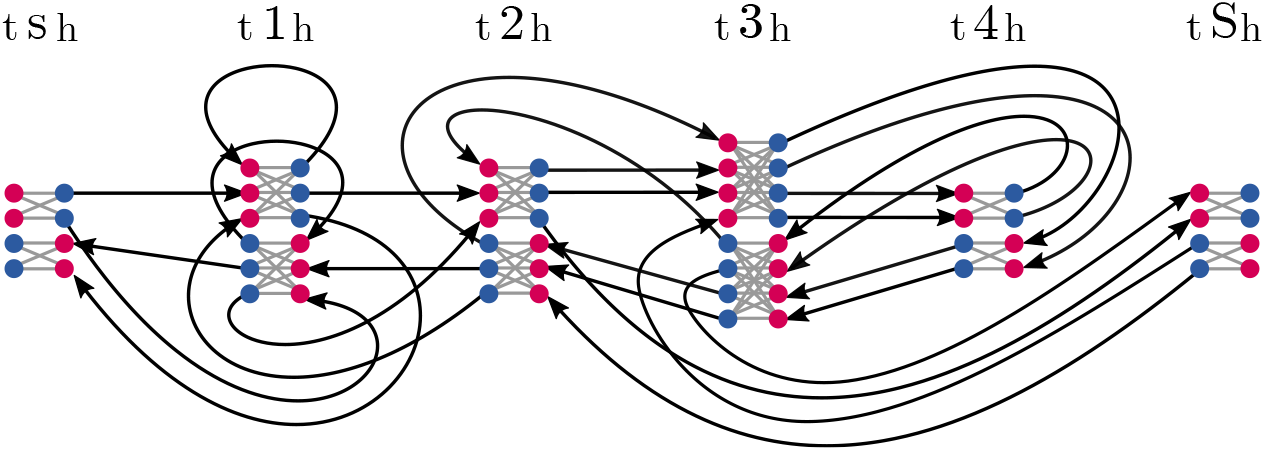

*In nodes and out nodes are highlighted in red and blue, respectively. For clarity, the direction of marker edges (gray edges; directed from in to out node) is omitted in the illustration*.

#### Graph decomposition

A perfect matching of marker edges in flow graph 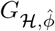 produces a set of alternating walks and alternating cycles through 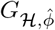, yet only half of the graph is eligible to form a solution to Problem 2. More precisely, for each marker *m* ∈ ℳ, exactly half of the number of its associated nodes in 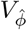 must be saturated in the matching that we seek, the other half as well as their incident edges must remain unsaturated. Further, we aim to admit only matchings that consist entirely of alternating 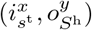-walks, because only those correspond to valid haplotypes of span(ℋ).

At last, we aim to assign to each saturated node 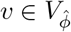 a position in some haplotype *A* of given haplotype set ℋ That way, we are able to determine whether the incident adjacency edge serves as continuation of the associated segment in *A*, or whether the incident saturated marker edge implies a recombination between two distinct segments.

The *integer linear program* (ILP) shown in Algorithm 1 implements the above-stated constraints.

#### Matching constraints

Each edge and node of flow graph 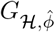 is associated with binary variables of x and y, respectively, that determine their saturation in a solution (cf. domains D.1 and D.2). Constraint C.01 ensures that each saturated marker edge is incident to saturated nodes. Perfect matching constraints, i.e., constraints that impose each saturated node being incident to exactly one marker edge, are implemented by constraint C.02. Similarly, constraint C.03 ensures that an adjacency edge is saturated iff its incident nodes are saturated. In other words, constraints C.01-C.03 together ensure that each component of the saturated graph corresponds to an alternating path or cycle component (the latter being prohibited by further constraints). The following two constraints C.04 and C.05 control the overall size of the saturated graph. In doing so, they ensure that, in a solution to Problem 2, the number of saturated nodes and adjacency edges matches the postulated multiplicity of markers *μ*(*m*), 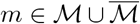, and pairs of consecutive markers 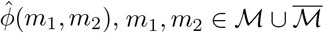, respectively.

#### Path constraints

Constraints C.05-C.08 force each component of the saturated graph to start and end in nodes associated with source *s*^t^ and sink *S*^h^, respectively, thereby ruling out any cycles. To this end, they make use of a set of integer variables `f` (cf. Domain D.03) that define an increasing flow within each saturated component that is bounded by constant *T* corresponding to the total flow of the graph, i.e., *T* := Σ_*m*ℱ ∈Σ_ *μ*(*m*). In each saturated marker edge, the flow is increased by 1 while along each adjacency edge, flow is kept constant. This prevents the formation of saturated cycles, because their flow would be infinite. Lastly, constraint C.08 preclude paths from starting in *S*^h^ or ending in *s*^t^, leaving only one option for any saturated component open, that is, the formation of a (*s*^t^, *S*^h^)-path.

#### Haplotype assignment

Each node in a solution to the ILP is associated with exactly one position in a haplotype in ℋ, recorded by binary variables `c`. Moreover, any marker edge whose incident pair of nodes is associated with the same position of the same haplotype corresponds to a conserved segment, i.e, no recombination within this marker has taken place. Each marker edge corresponding to a conserved segment contributes a score unit to the objective function. These score units are encoded by binary variables `t` (cf. domain D.05). Constraint C.09 ensures that each marker is associated with exactly one position *j* in a haplotype *A* of set 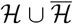, while C.10 confines incident nodes of adjacency edges to represent a consecutive marker pair *A*[*j*..*j* + 1]. At last, constraint C.11 allows t variables of marker edges to take on value 1 only if that marker edge is saturated and its incident nodes are associated with the same haplotype position.

By maximizing the sum over t variables, the objective minimizes the total number of segments needed to decompose the calculated founder sequences into segments from haplotypes 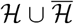 that are delimited by recombination events.

**Example 2** (cont’d). *The following plot illustrates a matching that is solution to Algorithm 1 for* 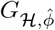. *The founder sequences are spelled out on the bottom, colored by haplotype (red, blue and green for haplotypes 2, 3 and 4 respectively). Unsaturated nodes and edges are grayed out, haplotype assignments implied by colored paths. The solution features two recombinations, marked by “*⋆*” along their associated marker edges*.

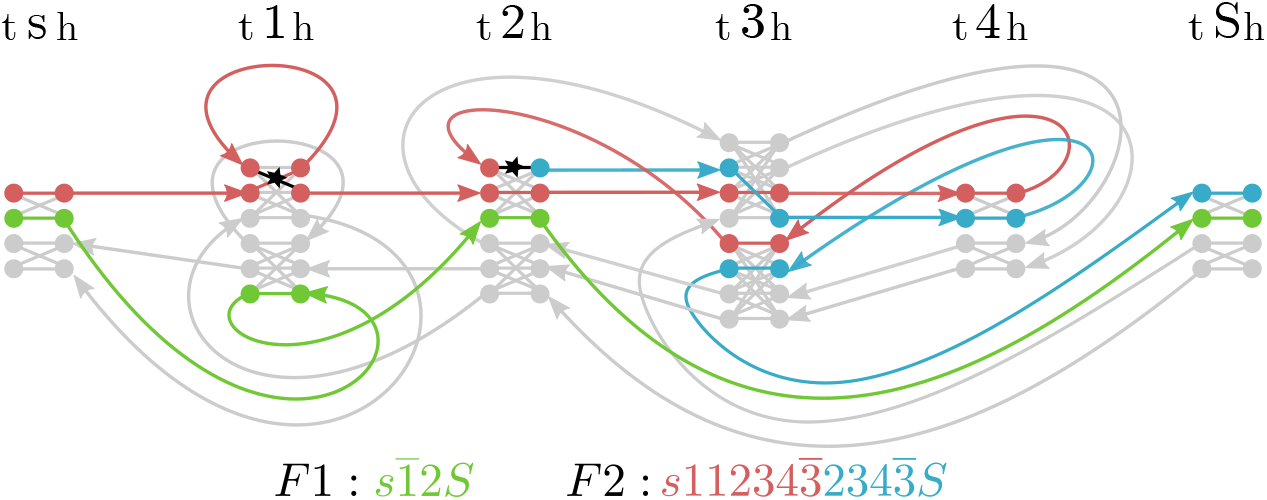

##### Algorithm 1

An ILP solution to Problem 2.

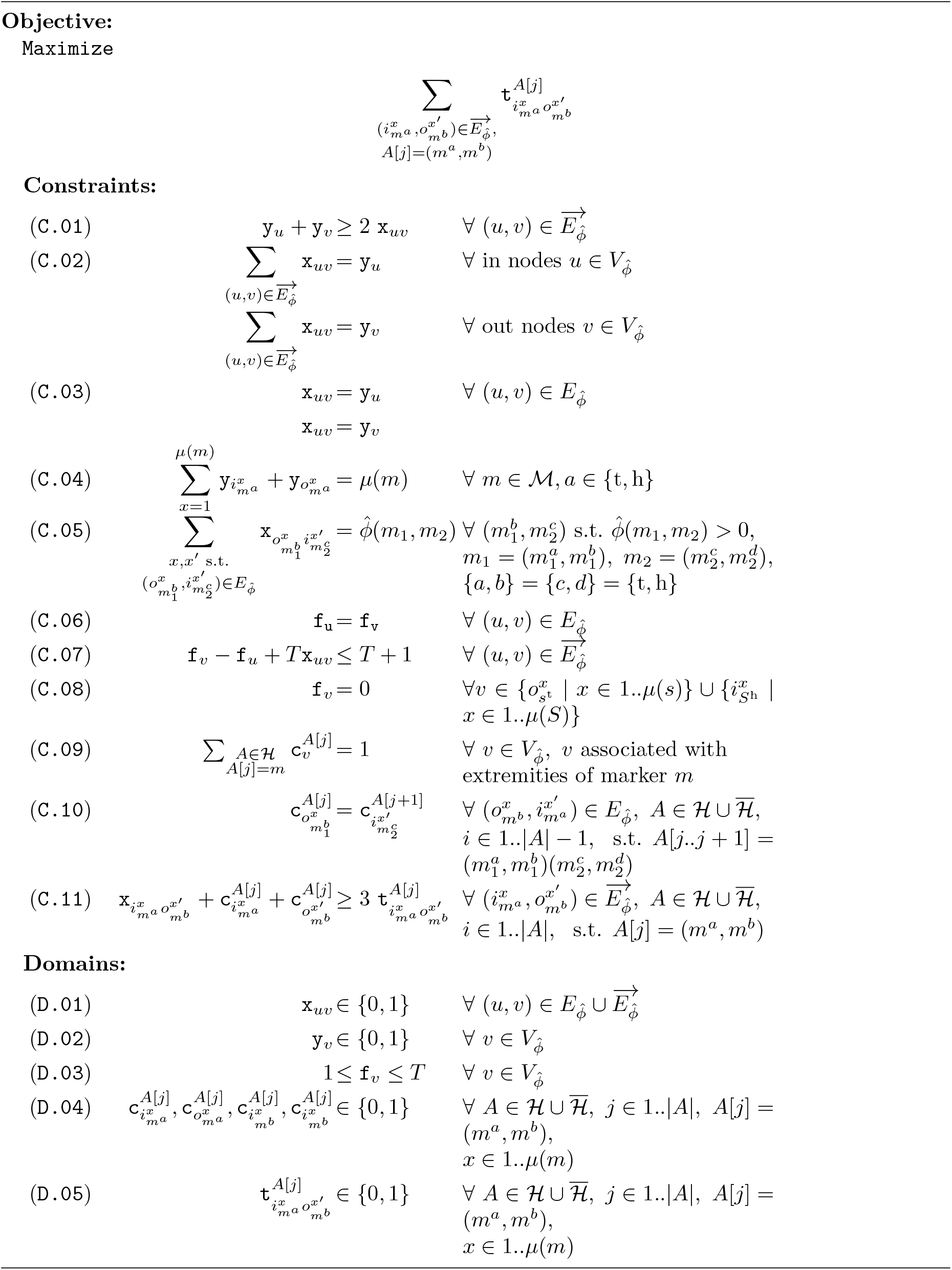

## 3 Results

We implemented our methods in the programming language Rust [14] and used Gurobi [11] as the solver. Our software is *open source* and publicly available on https://github.com/marschall-lab/hrfs. To run Algorithm 1 on a given set of haplotypes ℋ, we estimated the overall multiplicity 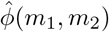 of pairs of consecutive markers *m*_1_*m*_2_ from a network flow solution to Problem 1 on ℋ. Note that this dispenses Algorithm 1 from being exact in our applications.

All experiments were run on a de.NBI cloud computing machine. For benchmarking purposes, we ran Gurobi single-threaded and recorded wall clock time (in seconds) and *proportional set size* (PSS, in Mb) for memory usage. Optimization time was capped at 30 minutes, beyond which the solver must capitulate and return its best-effort solution found thus far. The threshold for execution time is based upon available compute resources.

### 3.1 Experimental Data

We benchmarked the performance of our algorithms by conducting experiments on both simulated data and a real-world data set. The former presumed a simulator, capable of generating haplotypes with duplicated and inverted markers that can produce intricate homologous recombinations while providing control over the degree of complexity. To this end, we implemented our own simulation tool, that constructs a single haplotype sequence sampled at random to serve as seed. This seed sequence is adjustable by the following parameters: (i) number of distinct markers, i.e., the size of its variation graph, (ii) ratio of duplications, i.e., the number of additional edges inducing duplications in a walk of the graph, (iii) ratio of inversions, i.e., the proportion of inverted orientations within the set of duplications, and lastly (iv) the number of haplotypes that are input to subsequent founder set reconstruction. The latter are generated by performing random walks in the seed sequence’s variation graph and retaining only those leading from source to sink. In doing so, our simulator does not report nor have knowledge of a true founder set. Our simulator, discussed in more detail in Appendix A.1, enables us to explore various parameterizations that match different situations in biological data.

One important point concerns co-optimality. Problems 1 and 2 do not guarantee a unique solution. In fact, the pool of co-optimal solutions is often large for both problems. One contributing factor to co-optimality are cycles that are shared across multiple haplotypes, because they can be integrated in different orders. Further, the solution does not provide any information that could enable one to generate all co-optimal solutions nor discern between them, making a measure of accuracy challenging, since there is no guarantee that the “correct” founder sequence(s) will be seen in any number of trials.

In addition to simulated data, we applied our methods on a biological data set from the human 1p36.13 locus described by Porubsky *et al*. [21] to demonstrate their computational capabilities in realistic instances.

### 3.2 Simulation Experiments

To assess the impact of parameter configurations on the results, we ran a number of different experiments wherein all but one parameters are fixed. A reasonable choice of constants seemed to be 100 distinct markers, 10% of duplications, 10% of inversions and 10 haplotypes, motivated by our data on the 1p36.13 locus (8 markers, 68 haplotypes, 57% of duplications) and statistics compiled by Porubsky *et al*. [21] (6-7% duplications in the whole genome, <1% inversions).

#### Reduction in number of recombinations

To evaluate the efficacy of our solution to Problem 2, we compared the number of recombinations returned by Algorithm 1 to that in a solution obtained by our network flow algorithm for Problem 1. While the former is the immediate output of Algorithm 1, additional efforts needed to be made in order to retain the latter. In doing so, we estimate the number of recombinations in the flow solution by random assignment of corresponding segments in the original haplotype set and taking the one with the lowest number in 100k trials. Figure 2 summarizes the outcome of this experiment. Over all, Algorithm 1 found a solution with fewer recombinations in all instances but a few where Gurobi returned barely best-effort solutions after reaching the time limit of 30 minutes, all of which exhibited a gap of at least 100%. The parameter settings in those cases were extremal.

**Figure 2:**
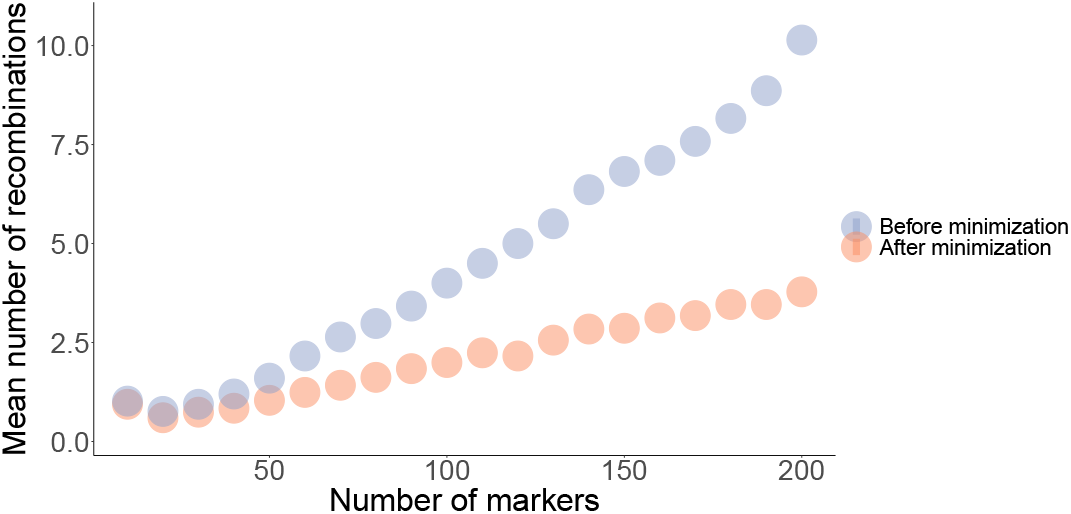
Mean number of recombinations by the size of the graph. Experiments were ran with values ranging from 10 to 200 in for the number of markers, in increments of 10. The ratio of duplications and of inversions was fixed to 10%, and number of haplotypes to 10. Each colored dot represents the mean number of recombinations over 50 replicates for one parameter set, after random assignment trials (blue) and after optimization (red).

Across all experiments and with a fixed ratio of duplications, inversions and number of haplotypes, the mean estimated number of recombinations both in the initial founder set and after minimization increases linearly with the number of markers, by approximately 4.2 and 2.0 per 100 markers respectively, reaching circa 10 and 3.8 for 200 markers. Results for experiments with other variable parameters are shown in Suppl. Figure S1.

#### Flow solution benchmark

Computing solutions with our network flow algorithm proved to be in almost all of our experiments near-instantaneous. By varying the number of distinct markers, the algorithm’s performance begins to deteriorate only with very large instances beyond 100k distinct markers and becomes excruciating for instances above 1M markers. When varying other parameters, we fixed the number of distinct markers to 100k rather than 100. Under 100k markers, execution completes after a mean wall clock time of 3.4 ± 2.0 seconds. In 95% of all experiments, the solver’s runtime was too short to make sufficient measurements for benchmarking memory usage; the maximum PSS for the remaining ones measured at 78MB. Over the 100k mark, both the graph size and duplication ratio begin to reduce performance, with an average runtime of 19.7 ± 8.7s. The ration of inversions on the other hand does not affect performance (Suppl. Figure S3). We measured peak memory consumption at 758MB across all conditions, which also occurred only at the very extremes of 100k distinct markers and a 100% ratio of duplications (Figure 3).

**Figure 3:**
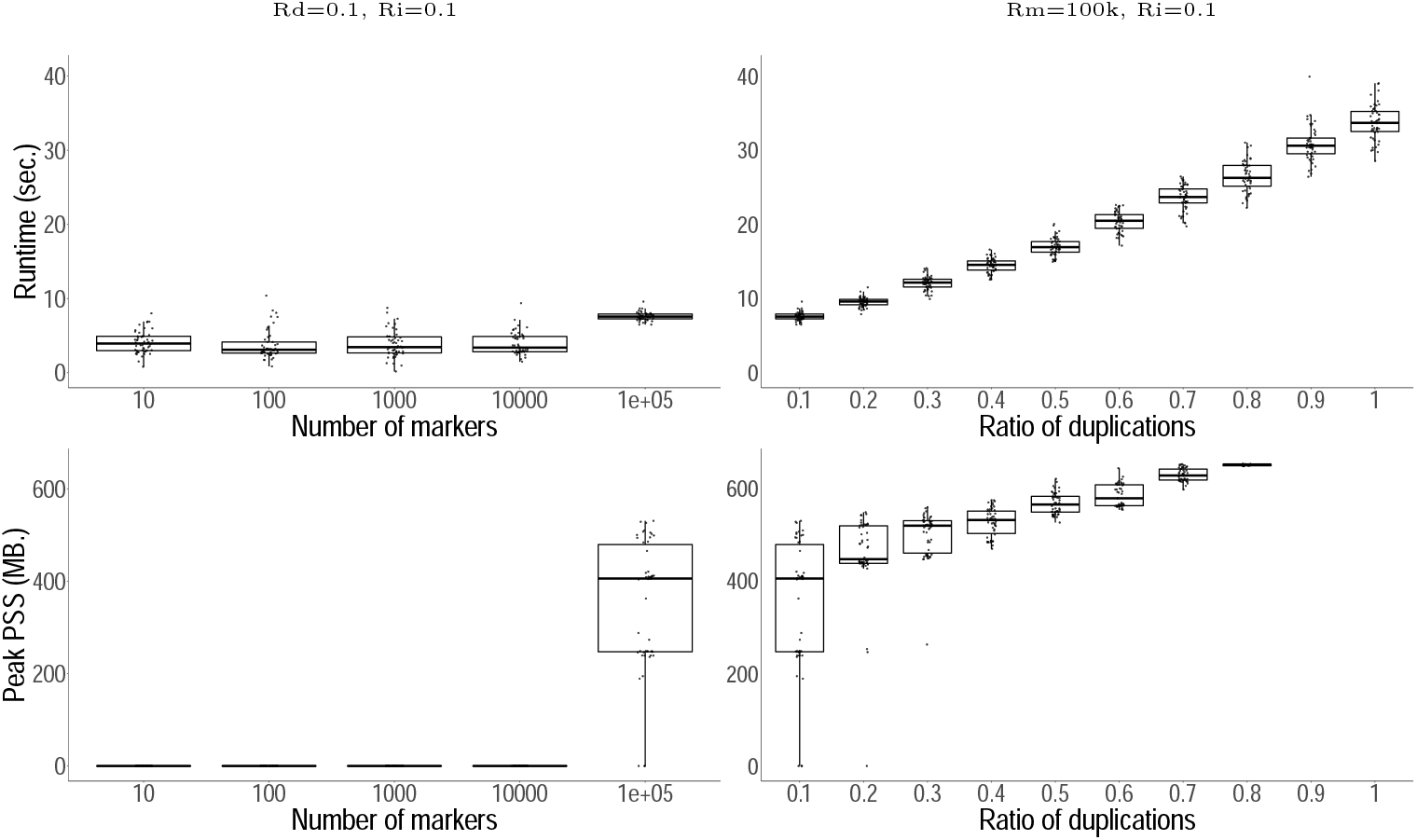
Problem 1, flow computational performance benchmarks. Runtime in seconds (upper panels) and peak PSS in Megabytes (lower panels), as a function of the number of markers (left) and of the ratio of duplications (right). For each experiment, the remaining parameters are fixed as indicated above. The abbreviations read as follows: *Nm*, number of markers; *Rd*, ratio of duplications; and *Ri*, ratio of inverted duplications.

#### Recombination minimization benchmark

As shown previously, Algorithm 1 successfully reduces the number of recombinations in solutions to Problem 1. However, its runtime increases dramatically with only moderate increments of any but one parameter of our simulator, the ratio of inversions; it does not play any role in performance (Suppl. Figure S2). For the remaining three, going beyond instances of 200 distinct markers, 20% of duplications, or 40 haplotypes typically does not allow for the optimization to finish in a reasonable amount of time (Figure 4, Suppl. Figure S2). A similar but much less pronounced trend is seen with memory usage, which still remains relatively low. Peak memory usage was again observed at extreme parameter values with a PSS of 1072MB with 50 haplotypes.

**Figure 4:**
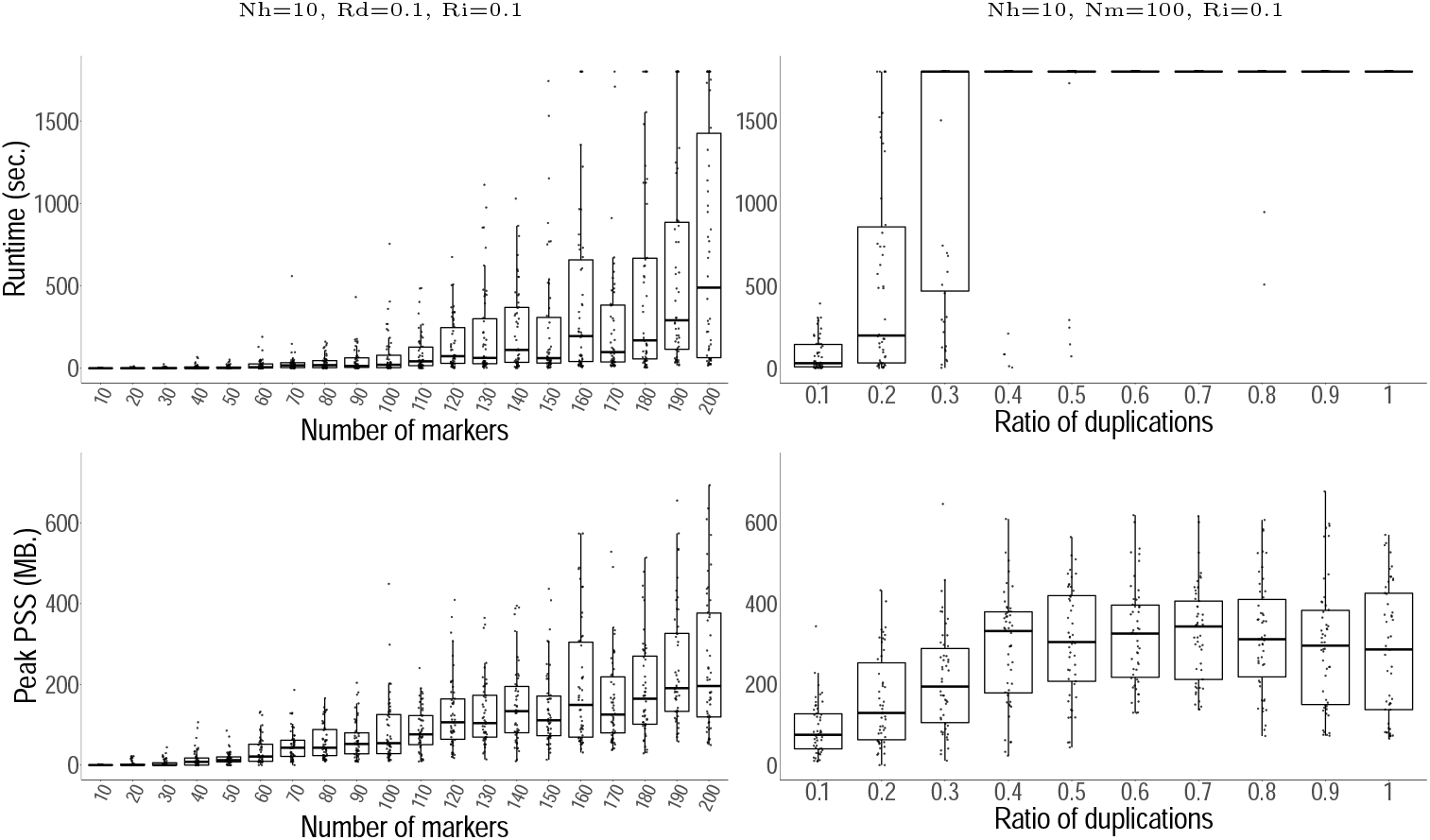
Problem 2, recombinations minimization performance benchmarks. Plots analogous to Figure 3. Runtime in seconds (upper panels) and peak PSS in Megabytes (lower panels), as a function of the number of markers (left) and of the ratio of duplications (right). For each experiment, the remaining parameters are fixed as indicated above. The abbreviations read as follows: *Nh*, number of haplotypes; *Nm*, number of markers; *Rd*, ratio of duplications; and *Ri*, ratio of inverted duplications.

### 3.3 Application: Locus 1p36.13

We obtained data from 68 human haplotypes (two per 34 individuals) at the 1p36.13 locus from Porubsky *et al*. [21] and the T2T-CHM13 human reference sequence [18]. The sequences comprise only eight distinct markers, terminal markers included. The sequences are attributed to five super populations, out of which 18 are of African origin (AFR), 16 of Eastern Asian (EAS), 12 of Admixed American (AMR), 12 of European (EUR), and 10 are South Asian (SAS). Their variation graph is densely connected with 26 edges (Figure 5). The 68 haplotypes display a high degree of genetic diversity, with haplotype sequences differing in order, orientation, and copy number of the marker (Suppl. Table T1). Haplotype lengths in terms of the number of markers vary from 15 to 26, with a median of 19.

**Figure 5:**
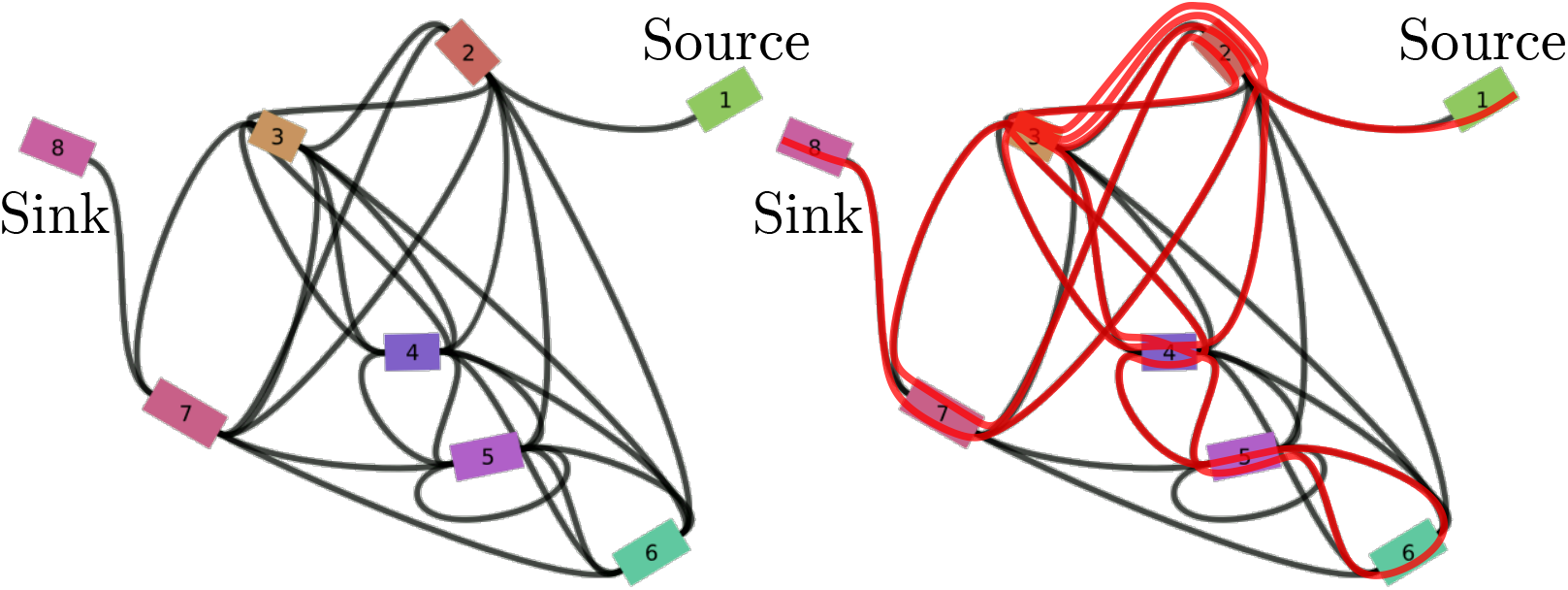
Graphical representation of the variation graph for the 1p36.13 locus data. On the left, a 2D plot rendered by Bandage [34]. Markers are represented as numbered colored rectangles, and the undirected edges connecting them as black curves. Markers 1 and 8 correspond respectively to the source and the sink of the graph. The right plot shows the walk through the graph corresponding to the sequence of haplotype `AFR-NA19036-h1`, a sample of African origin from our experimental data. The sample’s sequence in the previously established notation is: 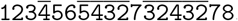.

Our network flow algorithm determined that the data set can be generated from a single founder sequence. Our randomized algorithm for calculation of the minimum number of recombinations in a solution to Problem 1 asserted 15 recombinations after 1M trials, while Algorithm 1 obtained an optimal solution that revealed only 9 recombinations. Minimization completed in 60.3 seconds with a peak PSS of 225MB. Note that there exists multiple other co-optimal solutions; Suppl. Figure S4 is an illustration of one.

## 4 Conclusion

The advent of sequencing technology and genome assembly methodology to reconstruct full human genomes enables research into previously inaccessible segmental duplication loci. This exciting opportunity entails a demand for explanatory models that can infer evolutionary relationships and histories of complex repetitive genomic regions. In this work, we propose a model capable of explaining a broad range of balanced and unbalanced genome rearrangements. Our experiments on simulated data and on the 1p36.13 locus demonstrate that our algorithmic solutions to the founder set problem and the problem of minimizing recombinations in founder sets are capable of processing realistic instances.

Importantly, the model we are proposing is based on a molecular mechanism with a well-established role in shaping segmental duplication architecture. In our view, many past models of genome rearrangements have not sufficiently captured biological reality and there is an important need for further research aiming to incorporate knowledge of molecular mechanisms into such models. For instance, we envision future models that additionally include mechanisms like non-homologous end joining (NHEJ) and mobile element insertions. Furthermore, actual rates at which NAHR occurs depend on factors like the length of the duplicated sequence, the sequence similarity, as well as the presence of specific sequence motifs. “Hidden” in our current approach in the construction of the variation graph, we aim to address and model these factors explicitly in future work.

## 5 Acknowledgements

The authors kindly thank Feyza Yilmaz for providing the haplotype data of the 1p36.13 locus.

## A Appendix

### A.1 Methods for the Simulation Experiments

The simulation experiments were carried out with the help of a new tool developed specifically for it. It generates a single *seed* haplotype, which then serves to construct a variation graph, on which random walks from source to sink are made to generate new haplotypes. The seed haplotype is initially a sequence of unique markers. A rate of duplications determines the number of duplications to add. For each duplication, the marker to duplicate and the position of insertion are sampled at random. The orientation of the duplicates is sampled according to a ratio of inversions. Next, the seed’s *variation graph* is built based on its sequence, represented as a walk through the graph. Finally, a given number of unique haplotypes is generated by performing random walks from source to sink in the graph. Essentially, the simulator starts from seed sequence, then generates an observable set of haplotypes and their graph. Because the walks are random, edges not covered by any of the new haplotypes must be pruned in order to respect the properties of a variation graph. The number of markers and haplotypes, and the ratios of duplication and inversion are the simulation parameters. The ratio of duplications (resp. of inversions) is defined as the ratio of the number of duplications to the number of nodes (resp. number of inversions to the number of duplications). All simulation experiments were carried out by running 50 simulations per parameter set, then applying the solutions of Problems 1 and 2 over the generated graph and haplotype set. The simulation experiments are implemented as *Snakemake* [15] workflows which also provide the benchmarking results then used for evaluation. The data and workflows for the 1p36.13 locus, as well as all simulation experiments are available in the github repository^7^ under the `examples` directory.

### A.2 Supplementary Figures and Tables

In the following figures, for each of the simulation experiments, performance is measured with regards to a range of values of a single parameter. All others are fixed to a constant value indicated above the given plot. They are labeled as follows: *Nm*, number of markers; *Nh*, number of haplotypes; *Rd*, ratio of duplications; and *Ri*, ratio of inverted duplications. Runtime is measured in seconds of wall clock time, and peak memory usage as the peak proportional set size (PSS) in Megabytes.

**Figure S1:**
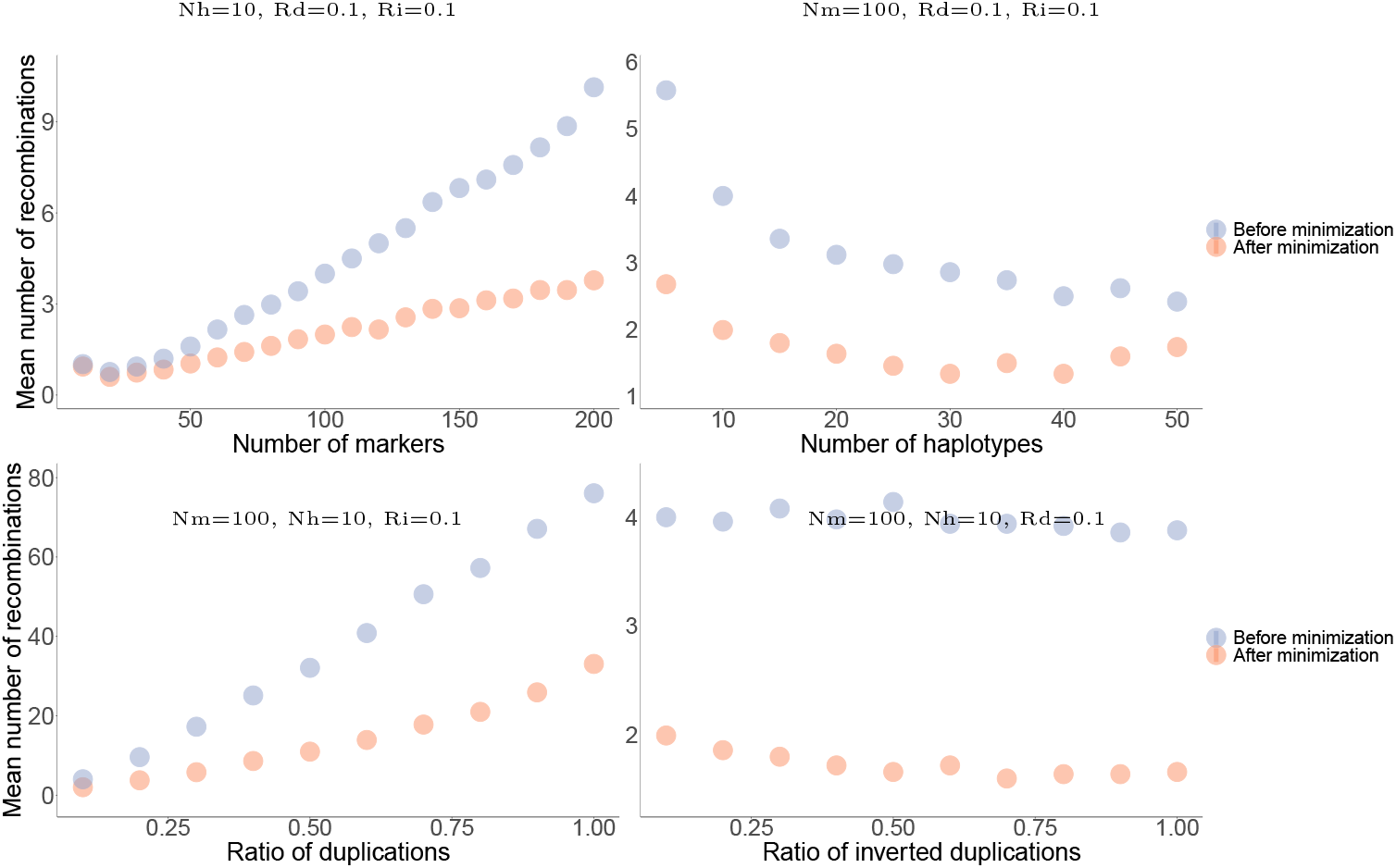
Reduction in the number of recombinations following minimization. The plots show the total number of recombinations before (blue dots) and after (red dots) minimization, as a function of each simulation parameter.

**Figure S2:**
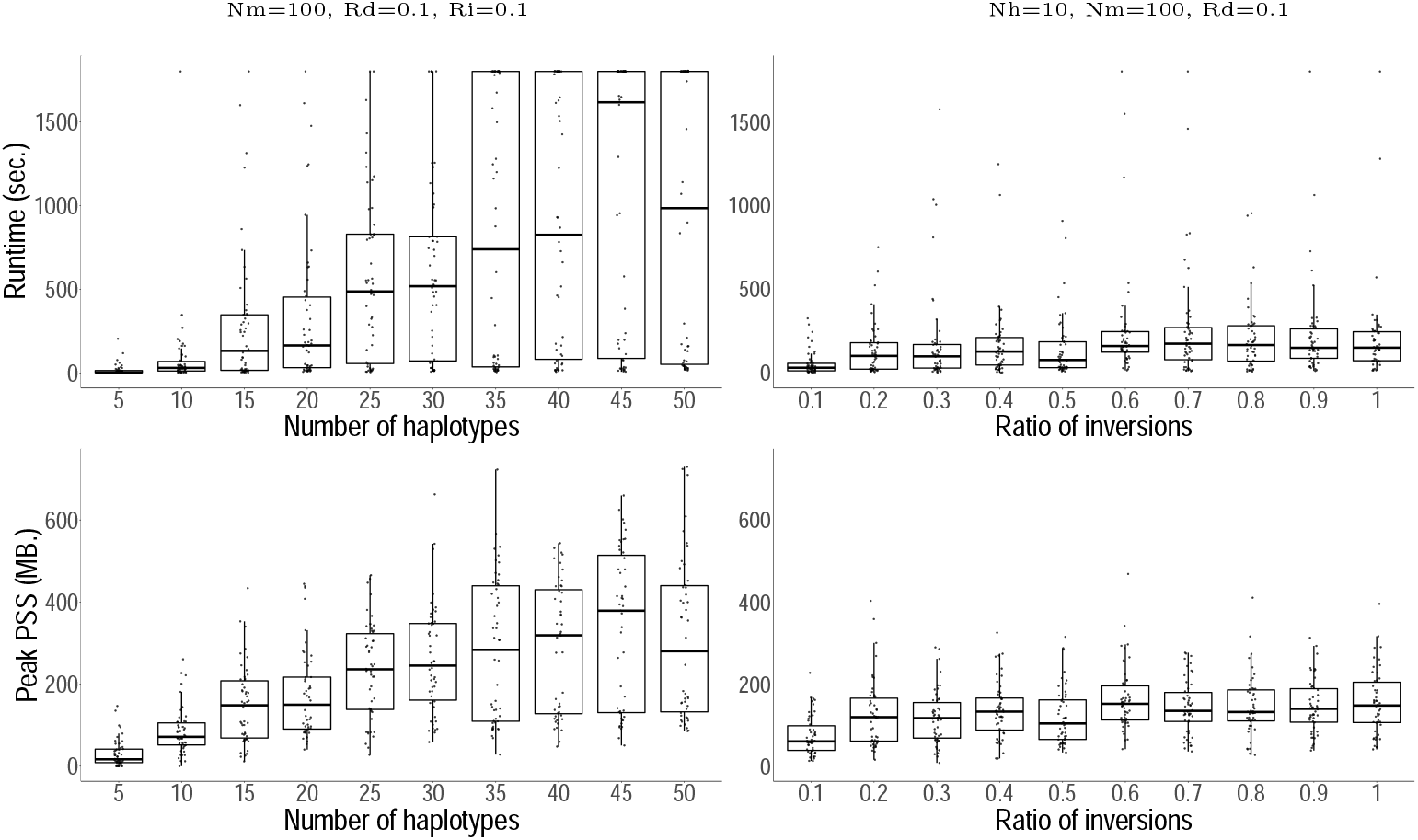
Number of recombinations minimization benchmarks. Runtime (upper panels) and peak PSS (lower panels) as a function of the number of haplotypes (left) and the ratio of inverted duplications (right).

**Figure S3:**
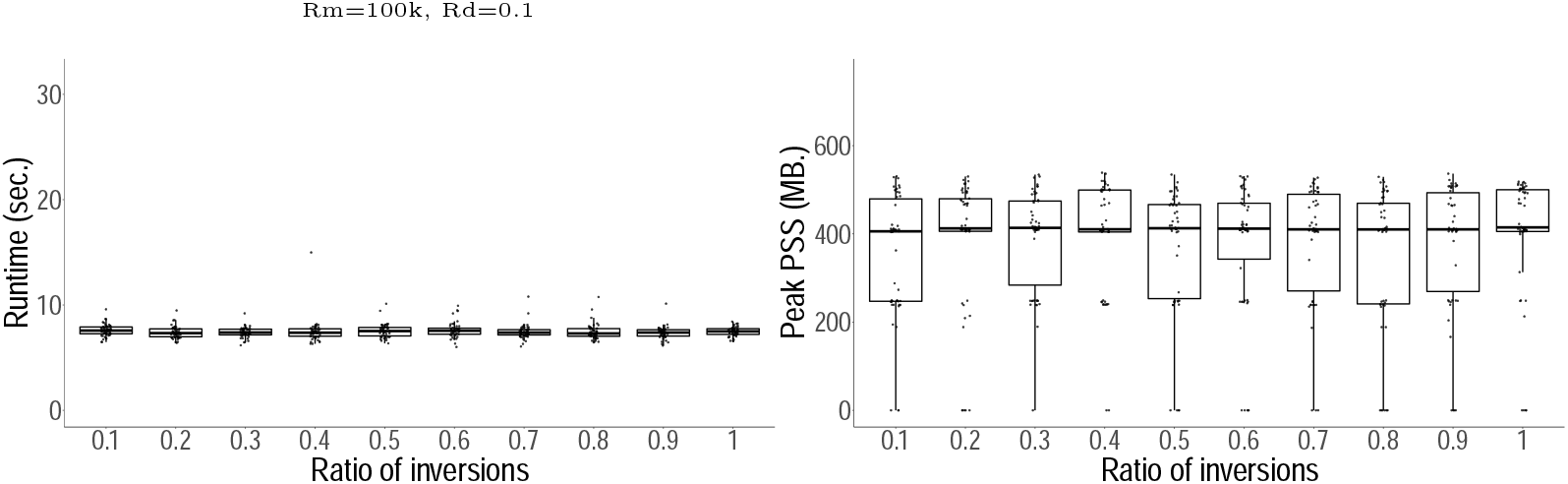
Flow computation performance with a variable ratio of inversions. Runtime (left) and memory usage (right) as a function of this parameter.

**Figure S4:**
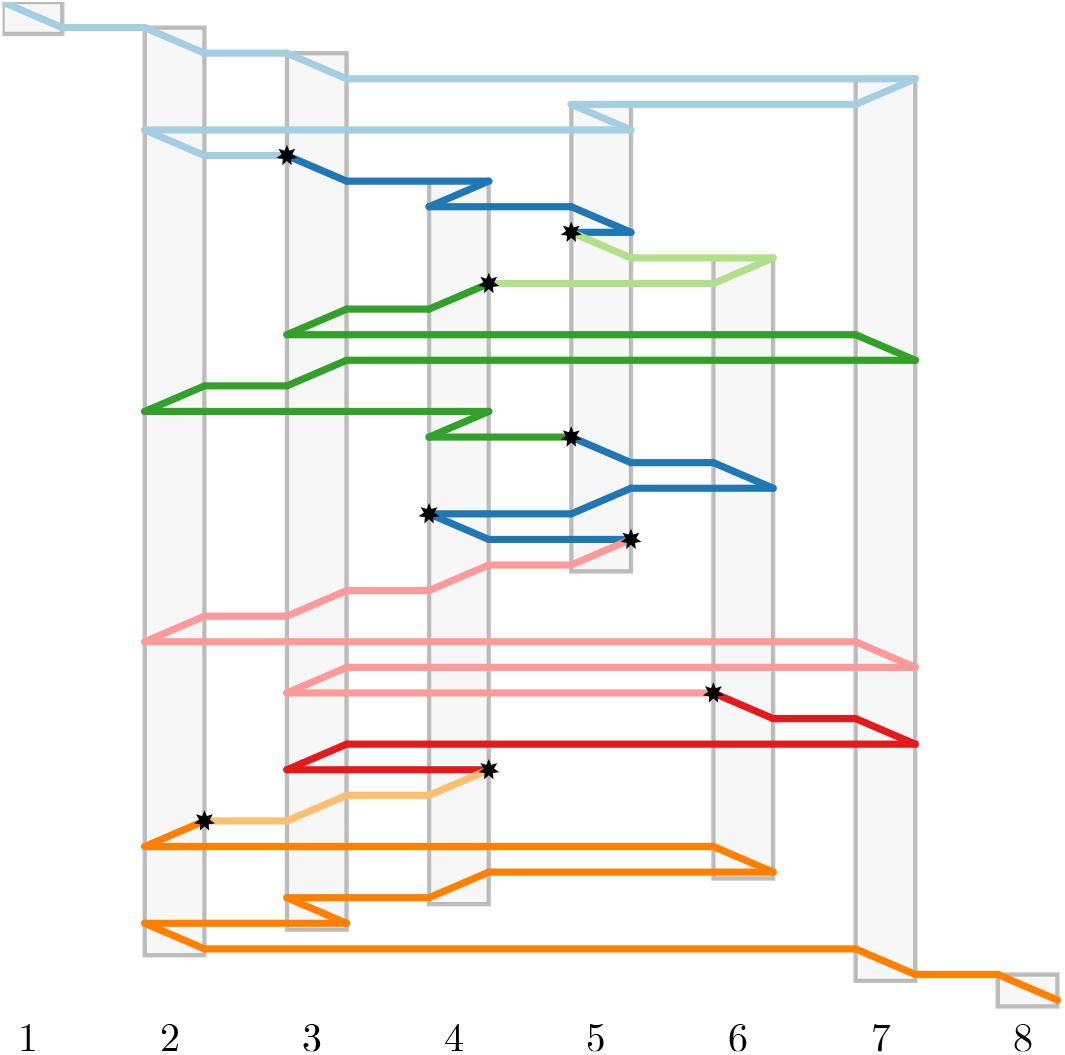
Visualization of a solution to the minimization problem on the 1p36.13 locus. The gray bars correspond to the graph’s nodes, labeled 1 to 8. The founder sequence `(>1>2>3<7>5>2>3<4>5>5<6<4<3>7<3<2<4>5>6<5>4<5<4<3<2>7<3>6>7<3<4<3<2>6<4>3>2>7>8)` is traced from top to bottom. A slanted line indicates the underlying node being traversed; if slanted rightwards, traversal is in forward direction, and if slanted leftwards, traversal is in reverse direction. Colors correspond to different haplotypes. The haplotype sequence is: *EUR-HG00171-h2, AFR-NA19036-h1, SAS-GM20847-h2, AFR-HG03065-h2, AFR-NA19036-h1, AFR-NA19036-h1, AMR-HG01573-h2, AFR-HG02011-h2, AFR-HG03371-h2, SAS-HG03683-h2*. Recombinations are marked with a star.

**Table T1:**
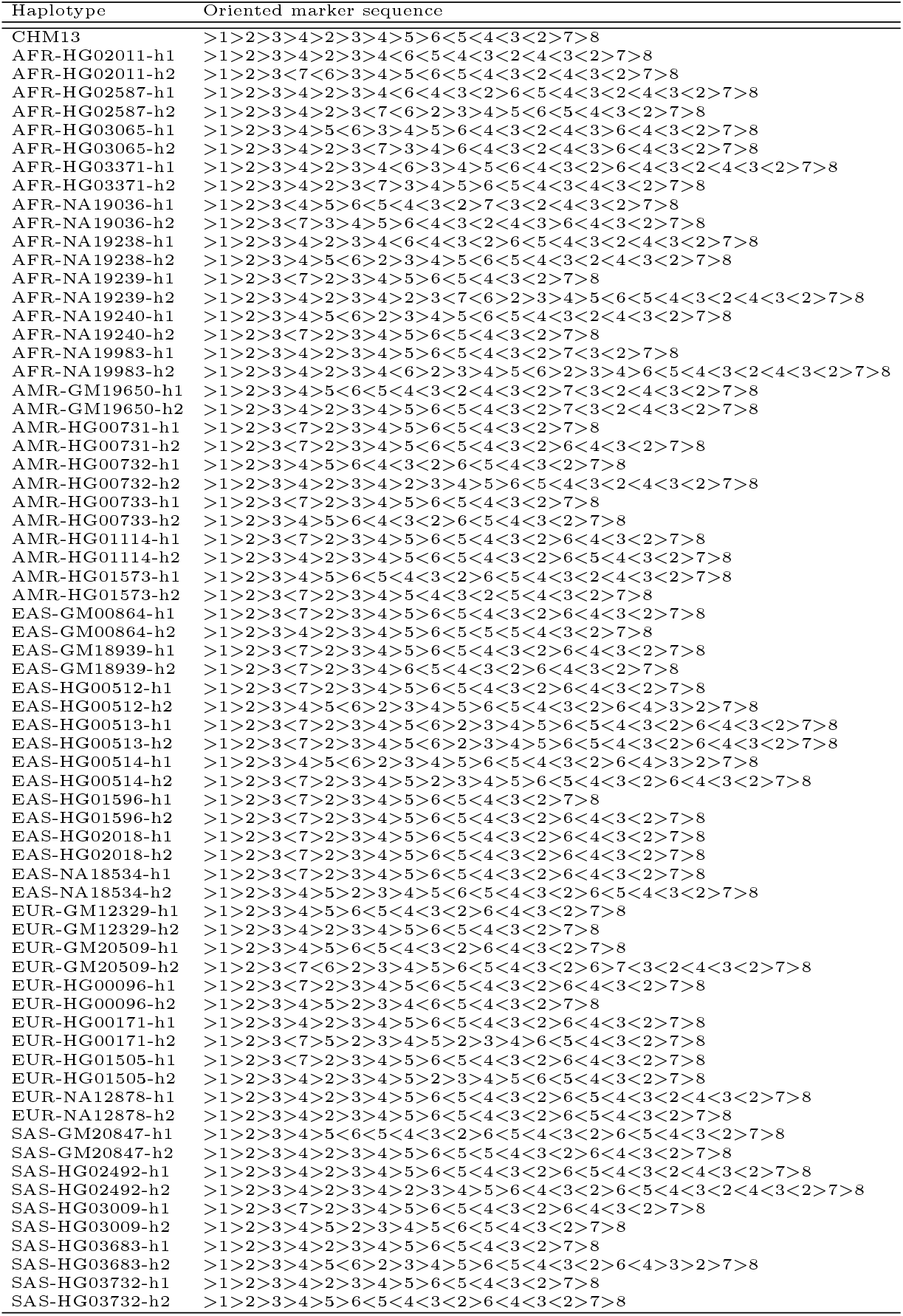
Haplotype marker sequences used in the 1p36.13 locus analysis, sorted alphabetically. The haplotype labeled *CHM13* is the provided reference. The sequences are in GFA Path format, where *>* corresponds to traversal in forward direction, and *<* in reverse direction.

## Footnotes

1 A version of this manuscript was accepted for publication at WABI 2022.

2 https://github.com/pangenome/pggb

3 Our notation is consistent with common practice of illustrating markers as arrows, that, in natural reading direction, face from left (tail of the arrow) to right (head of the arrow).

4 By construction, (*s*^t^, *S*^h^)-walks have out-degree 0, i.e., those with in-degree 0 are singleton in *G′*.

5 Our experiments directly use the results of Problem 1 as input for Problem 2. In other words, the multiplicities reported by 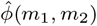 are the number of occurrences of (*m*_1_, *m*_2_) in a solution to Problem 1. This makes our experimental solutions to Problem 2 heuristic.

6 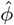 and *μ* are symmetric w.r.t. the relative orientation of markers, 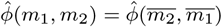 and 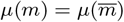

7 https://github.com/marschall-lab/hrfs

